# Pastures and Climate Extremes: Impacts of cool season warming and drought on the productivity of key pasture species in a field experiment

**DOI:** 10.1101/2020.12.21.423155

**Authors:** Amber C. Churchill, Haiyang Zhang, Kathryn J. Fuller, Burhan Amiji, Ian C. Anderson, Craig V. M. Barton, Yolima Carrillo, Karen L. M. Catunda, Manjunatha H. Chandregowda, Chioma Igwenagu, Vinod Jacob, Gil Won Kim, Catriona A. Macdonald, Belinda E. Medlyn, Ben D. Moore, Elise Pendall, Jonathan M. Plett, Alison K. Post, Jeff R. Powell, David T. Tissue, Mark G. Tjoelker, Sally A. Power

**Author notes:** Department of Ecology, Evolutionary Biology and Behavior University of Minnesota, St. Paul, MN, USA.

## Abstract

Shifts in the timing, intensity and/or frequency of climate extremes, such as severe drought and heatwaves, can generate sustained shifts in ecosystem function with important ecological and economic impacts for rangelands and managed pastures. The Pastures and Climate Extremes experiment (PACE) in Southeast Australia was designed to investigate the impacts of a severe winter/spring drought (60% rainfall reduction) and, for a subset of species, a factorial combination of drought and elevated temperature (ambient +3 °C) on pasture productivity. The experiment included nine common pasture and Australian rangeland species from three plant functional groups (C_3_ grasses, C_4_ grasses and legumes) planted in monoculture. Winter/spring drought resulted in productivity declines of 45% on average and up to 74% for the most affected species (*Digitaria eriantha*) during the 6-month treatment period, with eight of the nine species exhibiting significant yield reductions. Despite considerable variation in species’ sensitivity to drought, C_4_ grasses were more strongly affected by this treatment than C_3_ grasses or legumes. Warming also had negative effects on cool-season productivity, associated at least partially with exceedance of optimum growth temperatures in spring and indirect effects on soil water content. The combination of winter/spring drought and year-round warming resulted in the greatest yield reductions. We identified responses that were either additive such that there was only as significant warming effect under drought (*Festuca*), or less-than-additive, where there was no drought effect under warming (*Medicago*), compared to ambient plots. Results from this study highlight the sensitivity of diverse pasture species to increases in winter and spring drought severity similar to those predicted for this region, and that anticipated benefits of cool-season warming are unlikely to be realised. Overall, the substantial negative impacts on productivity suggest that future, warmer, drier climates will result in shortfalls in cool-season forage availability, with profound implications for the livestock industry and natural grazer communities.

## 1 Introduction

Climate change is a dominant driver of ecosystem change across the globe (Steffen et al., 2015; Sage, 2019). Exposure to high temperatures and changes in rainfall regimes have been shown to disrupt physiological function and alter plant species’ interactions, ultimately driving changes in productivity and ecological processes such as nutrient and water cycling (Parmesan et al., 2000; Backhaus et al., 2014; Frank et al., 2015; Ma et al., 2015). Predicting the impact of climate change on these processes is challenging, as temperature profiles (including minimum and maximum values) and the frequency, timing and size of rainfall events play major roles in driving change in ecosystem function, whereas climate models and projections often focus on only changes in mean annual temperature or precipitation (Easterling et al., 2000; Kreyling et al., 2008; Jentsch et al., 2009). Despite a recent increase in the number of studies focusing on climate extremes (De Boeck et al., 2015, 2019; Knapp et al., 2017; Hanson and Walker, 2019), relatively few have addressed the ecological implications of seasonal shifts in climate, which is important for understanding the underlying trade-offs between plant phenology and associated plant functional group responses (i.e. C_3_ vs. C_4_, legumes; Beier et al., 2012). This is especially true for studies considering the impact of multiple climate variables simultaneously as the combination may generate contrasting predictions for different plant functional groups (Beier et al., 2012).

Shifts in the seasonality of precipitation associated with climate change are predicted for many terrestrial ecosystems (IPCC, 2021). These changes are likely to have serious economic ramifications in biomes where productivity is tightly coupled with seasonal rainfall patterns (O’Mara, 2012; Godde et al., 2020), especially in warmer regions where the absence of low-temperature constraints allows year-round plant growth that depends on seasonal rainfall (Arredondo et al., 2016; Zeiter et al., 2016). Grassland phenology is often predicted based on classifications of plant traits or functional groups, with different optimum temperatures for photosynthesis (C_3_ vs. C_4_ grasses; Munson and Long, 2017; Ode and Tieszen, 1980; Winslow et al., 2003), timing of flowering in relation to peak summer temperatures (Sherry et al., 2007), life history strategies (annual vs. perennial; Cleland et al., 2006; Enloe et al., 2004; Veenendaal et al., 1996) and life forms (grasses/legumes/forbs etc.; König et al., 2018; Lesica and Kittelson, 2010; Rathcke and Lacey, 1985). In temperate and subtropical climates, grasslands dominated by C_3_ grasses mainly accrue biomass during cooler periods of the year (i.e. winter or spring), while C_4_-dominated grasslands typically accumulate biomass later in the growing season, when temperatures are higher (Pearcy and Ehleringer, 1984; Pearcy et al., 1987; Adams et al., 2016; Wilcox et al., 2016). These broad patterns are based on key differences in optimum temperatures for photosynthesis (Figure 1), as well as differences in water-use-efficiency and strategies for nutrient acquisition. Field-based manipulations examining impacts of drought on grassland production have primarily been conducted at the community scale, where both direct and indirect (inter-specific competition) responses combine to determine community-level productivity. For pasture systems, however, it is important to evaluate climate impacts for a wide range of forage species grown in monoculture in order to predict productivity responses and capacity to expand into new regions or vulnerability within current ranges as local climate shifts (Johnston, 1996; Bindi and Olesen, 2011).

**Figure 1.**
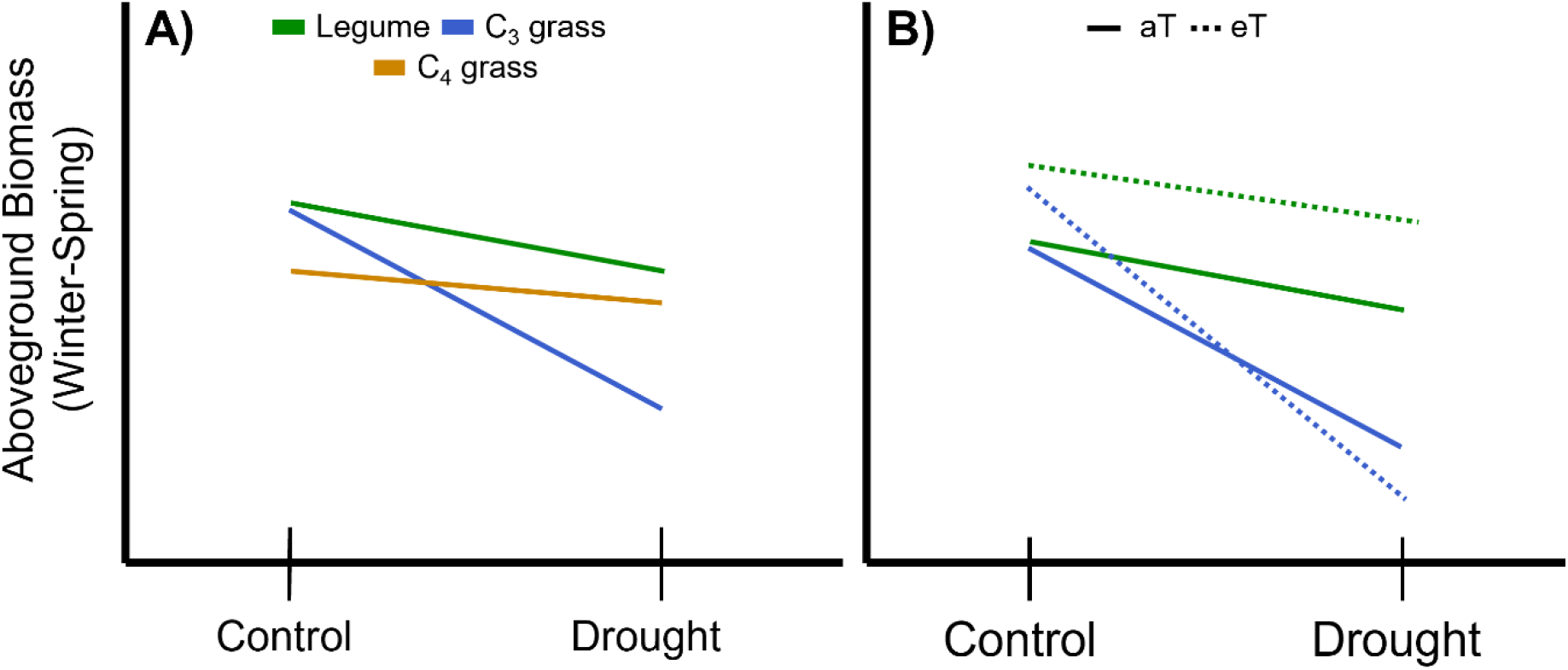
Schematic representation of expected changes in aboveground plant biomass during winter and spring associated with concurrent drought for A) plant functional groups including legumes and grasses relying on either the C_3_ or C_4_ photosynthetic pathways. Legumes are predicted to be less affected by drought (shallower slope) due to their higher nutrient and water use efficiency compared with C_3_ grasses, while C_3_ grasses are predicted to be most affected by drought due to a comparatively lower water use efficiency. C_4_ grasses are predicted to have lower biomass due to low temperature constraints on growth but lower drought sensitivity than C_3_ species due to greater water use efficiency. B) For legumes (green) and C_3_ grasses (blue) winter and spring warming (eT) is predicted to increase productivity under the control precipitation treatment relative to ambient temperature (aT; solid line); potential interactions between drought and warming can include an amplifying effect resulting in increased loss of biomass (C_3_ grass) or a stabilizing effect where warming offsets the impact of drought on biomass (legume).

In addition to changing rainfall regimes, rising temperatures are expected to have direct and indirect impacts on many ecosystem processes (Bai et al., 2013; Novem Auyeung et al., 2013; Song et al., 2019). In temperate climates, direct effects range from shifts in the timing of peak productivity relative to the start of the growing season (typically earlier) to extending the length of the growing season (Keatinge et al., 1998; Walker et al., 2006). The majority of field-based temperature manipulations have been conducted in temperate or cold-climate ecosystems, where low temperatures constrain growth for part of the year. These studies often find increased productivity and shifts in growth phenology, notably growth starting earlier and/or continuing later into the year, associated with warming (Aerts et al., 2006; Kueppers et al., 2017; Lu et al., 2013; Walker et al., 2006; Figure 1), although there are exceptions (Dukes et al., 2005; Arnone et al., 2011; Deutsch et al., 2011). Warming experiments in warm-temperate or subtropical climates with comparatively short mild winters and hot summers are relatively rare. These are, however, needed to evaluate the effects of rising temperatures in circumstances where warming results in exceedance of thermal optima for growth (Dukes et al., 2005; Volder et al., 2013; Song et al., 2019) and for regions where forage species grow throughout the year.

Plant functional groups are predicted to differ in their responses to warming, with warm-season species such as tropical grasses likely to benefit from increased winter or spring temperatures, while temperate grasses may be particularly vulnerable to higher spring or summer temperatures, especially when soil moisture is limiting (Munson and Long, 2017; Figure 1). Legume responses to warming are also likely to vary between seasons. For example, as with grasses, warming during the cooler months can bring temperatures closer to thermal optima (Sanz-Sáez et al., 2012; Whittington et al., 2013; Peng et al., 2020) ultimately promoting greater production (Figure 1). However, higher late spring and summer temperatures may result in growth reductions as a consequence of thermal constraints on nitrogen fixation and increased respiratory carbon losses or reduced photosynthesis if temperatures exceed temperature optima (Aranjuelo et al., 2007; Whittington et al., 2012). The impact of warmer temperatures on the timing and amount of biomass production in different species across seasons is an important knowledge gap, both in pastures and native grasslands and rangelands.

While studies that address the impact of either precipitation or temperature on plant physiology and ecosystem function are valuable, warmer temperatures frequently co-occur with drought (De Boeck et al., 2010; Dai, 2011; Yuan et al., 2016; Boer et al., 2020). Consequently, it is necessary to evaluate the impacts of co-occurring climate stressors on plant community structure and ecosystem function (Sherry et al., 2008; Hoeppner and Dukes, 2012). Climate models predict that temperate and sub-tropical Australia will be subject to large reductions in winter and spring rainfall (CSIRO, 2020) and grasslands across the globe are expected to experience more frequent and severe drought (Wang et al., 2021). Given the importance of pastures and rangelands for maintaining food security (O’Mara, 2012; Nábrádi, 2016; Godde et al., 2020), we established a large-scale, field experiment (at the **PA**stures and **C**limate **E**xtremes-PACE-field facility) to evaluate plant species’ responses to winter/spring rainfall reduction and year-round warming. Specifically, we asked: 1) How does winter/spring drought affect productivity in a range of pasture species? 2) Do drought responses in pasture grasses depend on their functional group? 3) Does cool-season warming enhance productivity and/or exacerbate the impacts of drought in the considered species?

## 2 Methods

### 2.1 Site description

The PACE facility was constructed in 2017 at the Hawkesbury Campus of Western Sydney University, in Richmond, New South Wales, Australia (S33.60972, E150.73833, elevation 25 m). Mean annual precipitation at this location is 800 mm (Australian Government Bureau of Meteorology, Richmond - UWS Hawkesbury Station 1980-2010); however, there is large inter-annual variability (annual precipitation 500-1400 mm over the past 30 years). Winter and spring precipitation accounts for 40% of annual rainfall. Mean annual temperature is 17.2°C, with the monthly maximum and minimum occurring in January (22.9° C) and July (10.2° C), respectively (BOM, 2020). The site is fenced to prevent access by mammalian herbivores. The soil is a loamy sand with a volumetric water holding capacity of 15-20% (**Table S1**).

The field facility comprises six replicate polytunnel shelters constructed from galvanized steel frames. These are covered with a single layer of 180 µm polyethylene (Argosee, Australia) to intercept all ambient precipitation, although the long sides are open to a height of 1.5 m to allow free flow of air (Figure 1). Shelters are 48 m long by 8 m wide, with a maximum height of 4.6 m, and are oriented along a SW-NE axis with the open ends facing into the direction of the prevailing wind. Each shelter has eight treatment plots (4 m by 4 m; Figure 1) that are further subdivided into four subplots (Figure 1B**; Figure S1**), with different plant species assigned at the subplot level (total 192 subplots). All surface soils were rotary-tilled to a depth of 12 cm to homogenize the upper soil profile prior to pasture establishment. All 4 x 4 m plots have a full root barrier installed to a depth of 90 cm to ensure hydrological isolation between treatments; the nested 2 x 2 m subplots have an additional root barrier between them to a depth of 30 cm to minimise root ingress.

Nine plant species were grown in monoculture subplots, along with three sets of two-species mixtures, for a total of twelve different planting combinations replicated in six independent shelters. However, for the purposes of this study, we focused on the nine monoculture species only. Species encompassed a range of functional diversity (C_3_ and C_4_ grasses, legumes; annuals and perennials) and origins (native grasses, and tropical or temperate introduced pasture grasses and legumes; Table 1) that are commonly found in improved grasslands (pastures) or rangelands (Clements et al., 2003). Sward establishment was initiated during early spring 2017 and yield data for the pilot year (2018-2019) are included in the supplement (**Table S2**). Initial sowing included a fertilizer addition in the form of diammonium phosphate (110 kg ha^-1^), and swards were subsequently managed via hand-weeding, herbicide and, where needed, insecticide application to maintain target species, in line with industry practice. Subplots with legumes received appropriate rhizobium inoculant during sward establishment: ALOSCA granular inoculant for *Biserrula* subplots (Group BS; ALOSCA Technologies, Western Australia, Australia); Easy Rhiz soluble legume inoculant and protecting agent for *Medicago* subplots (Group AL; New Edge Microbials, New South Wales, Australia). Subplots received top-up fertilization seasonally to replace nutrients removed from the soil (55 kg ha^-1^; Cal-Gran Aftergraze, Incitec Pivot Fertilisers, Australia).

**Table 1.** The origin, growth form and photosynthetic pathway of pasture and rangeland species selected for study in a drought and warming field experiment in the PACE facility

| Species* | Origin | Growth Form | Photosynthetic pathway | Warming Treatment |
| --- | --- | --- | --- | --- |
| *Biserrula pelecinus* | Temperate, introduced | Legume | C<sub>3</sub> |  |
| *Medicago sativa* | Temperate, introduced | Legume | C<sub>3</sub> | Yes |
| *Festuca arundinacea* | Temperate, introduced | Grass | C<sub>3</sub> | Yes |
| *Lolium perenne* | Temperate, introduced | Grass | C<sub>3</sub> |  |
| *Phalaris aquatica* | Temperate, introduced | Grass | C<sub>3</sub> |  |
| *Rytidosperma caespitosum* | Temperate, native | Grass | C<sub>3</sub> |  |
| *Chloris gayana* | Tropical, introduced | Grass | C<sub>4</sub> |  |
| *Digitaria eriantha* | Tropical, introduced | Grass | C<sub>4</sub> |  |
| *Themeda triandra* | Tropical, native | Grass | C<sub>4</sub> |  |

\*Species referenced by genus names in the text

### 2.2 Experimental treatments

All nine species were exposed to a winter/spring drought treatment, and a subset of two species (*Festuca*, *Medicago*) received a warming treatment in a factorial combination with drought (**Figure S1**). The drought treatment comprised a control (Control: C) and a drought (Drought: D) watering regime that was applied during the 6-month austral winter/spring period (1 June to 30 November 2019). A 60% reduction in winter/spring rainfall was chosen for the drought manipulation as representing the upper end of climate model predictions for end-of-century seasonal rainfall change for south-eastern Australia, under RCP8.5 (CSIRO, 2020). Furthermore, a 60% reduction in winter/spring rainfall aligns with historical climate extremes for key pasture growing regions across south-eastern Australia (BOM, 2019); this treatment therefore represented historically relevant rainfall extremes, which are predicted to increase in both frequency and duration. The control was set to represent a typical precipitation regime for the local area, accounting for long-term patterns in seasonality and in the statistical distribution of event sizes and timing within seasons (**Figure S2**).

The warming treatment comprised a year-round temperature increase of +3 °C (Table 1) achieved using infra-red (IR) heaters, approximating predicted changes in temperature for Australia by 2080 under RCP7.0 (IPCC, 2017) and SSP3-7.0 (IPCC, 2021) scenarios. Elevated temperatures (eT) were applied to two 4 x 4 m plots within each shelter; one plot received the control irrigation (eT-C), while the other plot received the drought treatment (eT-D) and these were paired with two ambient temperature (aT) plots, one receiving control irrigation (aT-C) and the other the drought treatment (aT-D). Each warmed plot had a heating array comprising eight 1000W ceramic heaters (FTE 1000W, Ceramicx, Ireland) mounted on an aluminium frame (4 m x 4 m) suspended 1.4 m above ground level (Figure 2C). Lamps were positioned to give uniform coverage of IR radiation across the four composite subplots. The power level to the heating lamps was adjusted each minute, via pulse width modulation using a solid-state relay controlled by a data logger (CR1000, Campbell Scientific), based on a proportional-integral control algorithm. Target temperatures for these plots were controlled via feedback from IR-sensors (SI-100, Apogee Instruments, Logan, UT, USA) mounted at a height of 3.8 m, recording plot surface temperatures every five minutes; temperatures thus represent plot-level means for the plant canopy and, where visible, soil, and are henceforth referred to as canopy temperatures. The + 3 °C warming treatment was applied with reference to canopy temperatures for the relevant control and drought treatments (i.e., aT-C paired to eT-C and aT-D paired to eT-D) to account for differences in soil moisture and vegetation cover between these plots.

**Figure 2.**
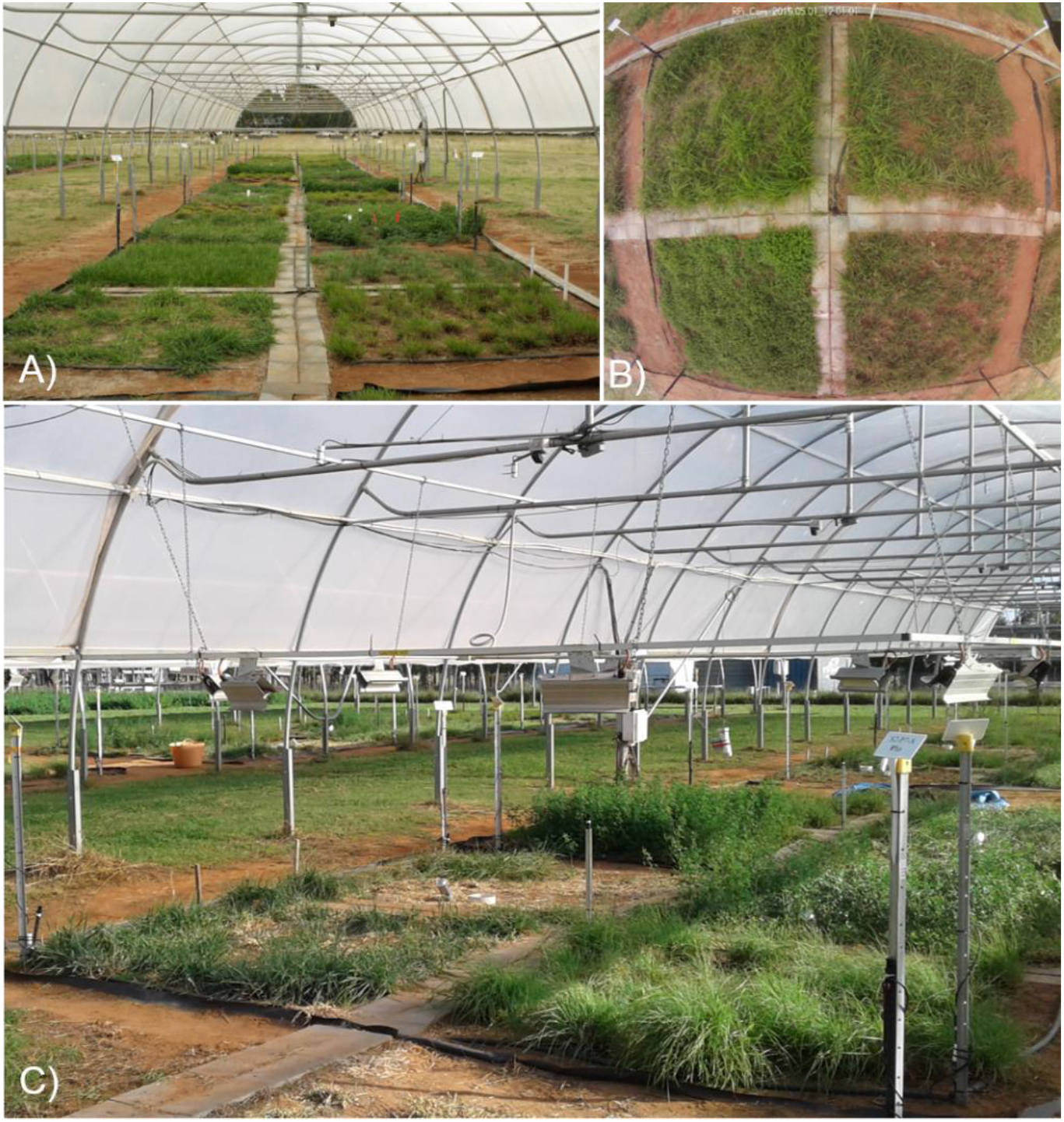
The Pastures and Climate Extremes (PACE) field facility located at Western Sydney University in Richmond, New South Wales, Australia. A) There are six open-sided polytunnels, each with eight experimental plots; B) Experimental plots (4 x 4 m) received a drought, warming, or drought + warming treatment and are each divided into four discrete 2 x 2 m subplots comprising nine different pasture species in monoculture; C) Heater (infra-red: IR) arrays were mounted above the vegetation canopy and warmed the plot surface an average of 3 °C above paired (control, drought) ambient temperature plots.

### 2.3 Environmental monitoring

Each shelter had a data logger (CR1000, Campbell Scientific) that recorded environmental conditions and regulated the heating lamps. Soil moisture sensors (16 per shelter; Time Domain Reflectometers; CS616, Campbell Scientific) recorded volumetric soil water content (0-15 cm) every 15 min in all six replicates of four different species subplots and treatment combinations (**Table S3**); in the *Medicago* subplots soil water content was also monitored at a second depth (15-30 cm). Soil temperature probes (T107, Campbell Scientific) were installed in the top 6-12 cm of the soil of eight subplots per shelter (all four drought and warming treatment combinations of *Festuca* and *Medicago*) to record soil temperature every 15 min (**Table S3**). Air temperature and humidity sensors (Series RHP-2O3B, Dwyer Instruments Inc, USA) mounted in force-ventilated radiation shields were installed inside three of the rainout shelters at 0.6 m height, with records collected every 5 min to determine any shelter effects on environmental conditions. Additionally, three sets of sensors were installed at the same height outside the shelters. Photosynthetically active radiation (PAR) was recorded at 5-min intervals using PAR sensors (Apogee quantum sensor, USA) installed at a 6 m height outside two shelters, with two additional sensors located within shelters at 3 m.

### 2.4 Biomass harvests

All subplots were regularly harvested by clipping to determine aboveground productivity during active growing periods. This ‘surrogate grazing’ involved use of hand shears and a sickle mower. Timing of harvests was based on grazing recommendations for individual species (Clements et al., 2003); hence, there were 2 (*Chloris* and *Digitaria*) or 3 (all other species) harvests per species during the six-month winter and spring period in 2019 (Clark et al., 2016). During each harvest plants were cut to 5 cm above the soil surface and weighed (fresh and dry mass), with a sub-sample of harvested material sorted to determine the proportion of live and dead biomass. The weed (i.e. non-target species) fraction from each subplot was also assessed and was excluded from aboveground biomass measurements (< 5%). All materials were oven-dried at 70 °C for at least 48 hours prior to determining dry mass.

### 2.5 Calculations and statistical analyses

The responses of temperature and soil water content to drought and warming treatments were determined based on subplot-level (soil moisture, soil temperature) or plot-level (canopy temperature) daily mean values. Aboveground production responses were determined based on subplot totals, summed across all harvests conducted during the 6-month winter/spring drought period.

Statistical analyses of treatment effects were conducted using linear mixed effects models with climate treatments (‘drought’ or ‘control’: D or C, ‘warming’ or ‘ambient’: eT or aT) as fixed effects and random effects defined as ‘subplot nested within plot’ or ‘plot’ (to account for non-independence among continuous measurements) nested within ‘shelter’ (to account for the blocked design), and ‘date’ (to account for temporal variation). Volumetric water content data in *Medicago* subplots were also analysed for differences between upper (0-15 cm) and lower (15-30 cm) depths among climate treatments by including ‘depth’ as a fixed effect. Climate treatment effects on aboveground production were analysed using two different linear mixed effects models based on our hypotheses comparing responses among different functional plant groups using a ‘functional group’ categorical predictor fixed effect. For the first set of these analyses, ‘drought’ and ‘warming’ treatments were included as fixed effects; ‘Species’ was also included as a random effect for models examining differences among ‘functional groups’ (as a fixed effect), and ‘plot nested within shelter’ was included as a nested random effect for all models. Aboveground production was natural log-transformed to meet assumptions of constant variance, where indicated in statistical tables. All analyses were conducted in R version 4.0.0 (R Core Team, 2020) using the package lme4 (Bates, 2015) and Kenward-Roger degrees of freedom were calculated using the Anova function in the ‘car’ package (Fox and Weisberg, 2019). Pairwise comparisons to determine treatment effects on soil or canopy temperature and soil water content, or aboveground production among species or functional groups, were conducted using the package ‘emmeans’ (Length, 2020) and he Tukey method for *P*-value adjustment.

To compare responses to drought among species and functional groups, log response ratios between paired control and droughted subplots within a shelter were calculated for aboveground biomass summed across all harvests to derive a treatment effect size:

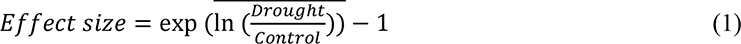

Linear mixed effects models were used for this second set of analyses evaluating differences in the effect size among 1) species and 2) functional groups, with both specified as fixed effects. Random effects included ‘shelter’ for both models, while the model for 2) also included ‘species’ as an independent random effect. Modelled data were back transformed prior to visualization of species-and functional group-level differences. The standard error of the mean effect size was calculated as the product of the back-transformed mean and the standard error of the effect size.

Interactions between warming and drought, the effect of drought on warming responses, and the effect of warming on drought effects, were calculated as the log response ratio between single factor treatment effects within a shelter for each period (Dieleman et al., 2012). These ratios were calculated such that the effect of drought was examined by comparing the effect size (Equation 1) for each warming level (aT-D / aT-C, eT-D / eT-C). The effect of warming was examined as the effect size for each drought level (eT-C / aT-C, eT-D / aT-D) and the effect of warming on the drought response as the ratio between these two effects ((eT-D / aT-D)/(eT-C / aT-C)).

## 3 Results

### 3.1 Drought and warming treatment effects on environmental conditions

Drought and warming significantly altered soil water content and temperature across the six-month winter/spring study period (Figure 3, **Figure S3-6**, **Tables S4** and **S5**). During this time, droughted subplots of all continuously monitored species (*Biserrula*, *Festuca*, *Medicago*, and *Lolium*) had significantly reduced soil moisture in the upper 15 cm (Figure 3; **Figure S3**; *P* < 0.05) compared with controls. Additionally, soil water contents in all droughted subplots were less variable compared to their respective control subplots (Figure 3A, B).

**Figure 3.**
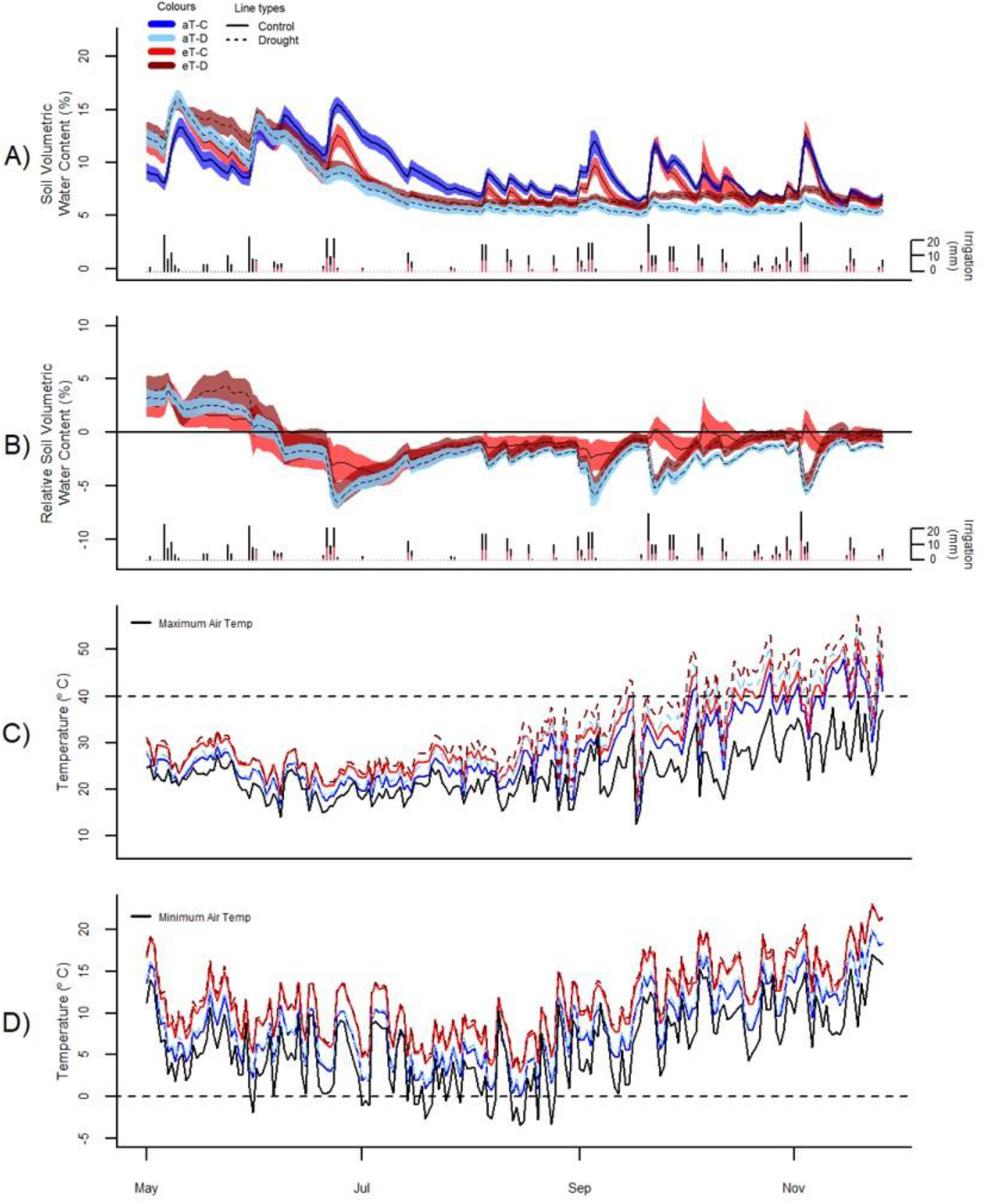
Experimental treatment effects of winter/spring drought (control: C, drought: D) applied during the period between 1 June to 30 November 2019 and year-round warming (ambient temperature: aT, elevated temperature: eT) treatments on soil moisture (panels A and B) and canopy temperature (panels C and D) averaged across the six shelters, from 1 May 2019 to 30 November 2019. A) Average soil volumetric water content in *Festuca* sub-plots (additional species are shown in Figure S2) with 95% confidence intervals as well as individual irrigation events as daily rainfall totals for control (black bars) and drought (red bars) plots over the 6-month period of study; B) Treatment differences in soil water content from aT-C plots over time in the *Festuca* subplots, noting the winter/spring drought treatment period between 1 June and 30 November 2019; C) Daily maximum canopy temperature, relative to maximum ambient air temperature (black line) and 40 °C (representing extreme temperatures, dashed lines); D) Daily minimum temperature compared with minimum air temperature (black line) and 0 °C (dashed lines).

Soil water content was reduced by warming in *Festuca* subplots, particularly under drought conditions (*P* = 0.03; **Figure S2**), although there were intermittent short-term positive effects of warming on soil moisture following irrigation events for this species (Figure 3B). Soil water content (0-15 cm) in *Medicago* subplots was unaffected by warming. There was, however, evidence of greater soil water content at depth (15-30 cm) in droughted *Medicago* soils, compared with surface levels (**Figure S4**).

Plot-level canopy temperatures were consistently increased by both drought (*P* < 0.001, + 1.0 °C) and warming (*P* < 0.001, + 3.0 °C). There were, however, no interactions between drought and warming due to the experimental design which referenced warming treatments to ambient temperatures for the respective droughted or control plot (**Figure S5**; **Table S4**). Additionally, warming altered the maximum and minimum temperatures and thus the temperature range within plots (**Figure S5**; **Table S5**). Warmed plots of *Festuca* and *Medicago* had no days when minimum canopy temperatures fell below freezing (compared to one day in ambient plots) and only 15 days when canopy temperatures fell below 5 °C (compared to 45 days in ambient plots). At the other end of the scale, the warming treatment resulted in an additional 16 days when canopy temperatures exceeded 40 °C and an extra 8 days where temperatures exceeded 45 °C (Figure 3C, D). Subsurface soil temperatures were also significantly increased by both drought (*Festuca*) and warming (*Festuca* and *Medicago*) (**Figure S6**). Overall, soil temperature responses were similar between species, although drought amplified the warming effect in *Festuca* but not *Medicago* soils.

Shelter effects on air temperature, relative air humidity, and PAR levels were even across shelters; on average there was an 11% decrease in temperature, a 6% increase in relative humidity and a 22% reduction in PAR under shelters, as compared to outside (**Figure S7**). Shelter effect on air temperature changed from winter to spring, with a slight cooling effect during the winter, likely due to lower levels of radiation, and gradually transitioned to a neutral and then positive effect during late spring (**Figure S7b**).

### 3.2 Aboveground production response to drought

During the 6-month winter/spring drought period, total aboveground production ranged from 2,800 to 8,300 kg ha^-1^ under control conditions (Figure 4A). The most productive species were *Themeda*, *Medicago* and *Rytidosperma*, while *Festuca*, *Chloris* and *Biserrula* were the least productive. Droughted subplots were significantly less productive than their respective controls, with an average yield reduction of ∼45% across the nine species (Figure 4B; Table 2). There was also a significant interaction between drought and species, with eight of the nine species having significantly lower productivity under drought (Table 2). The remaining species (*Lolium*, a perennial C_3_ grass) experienced late spring die back and had a non-significant 11% reduction in total productivity summed across the six months (*P* = 0.51).

**Figure 4.**
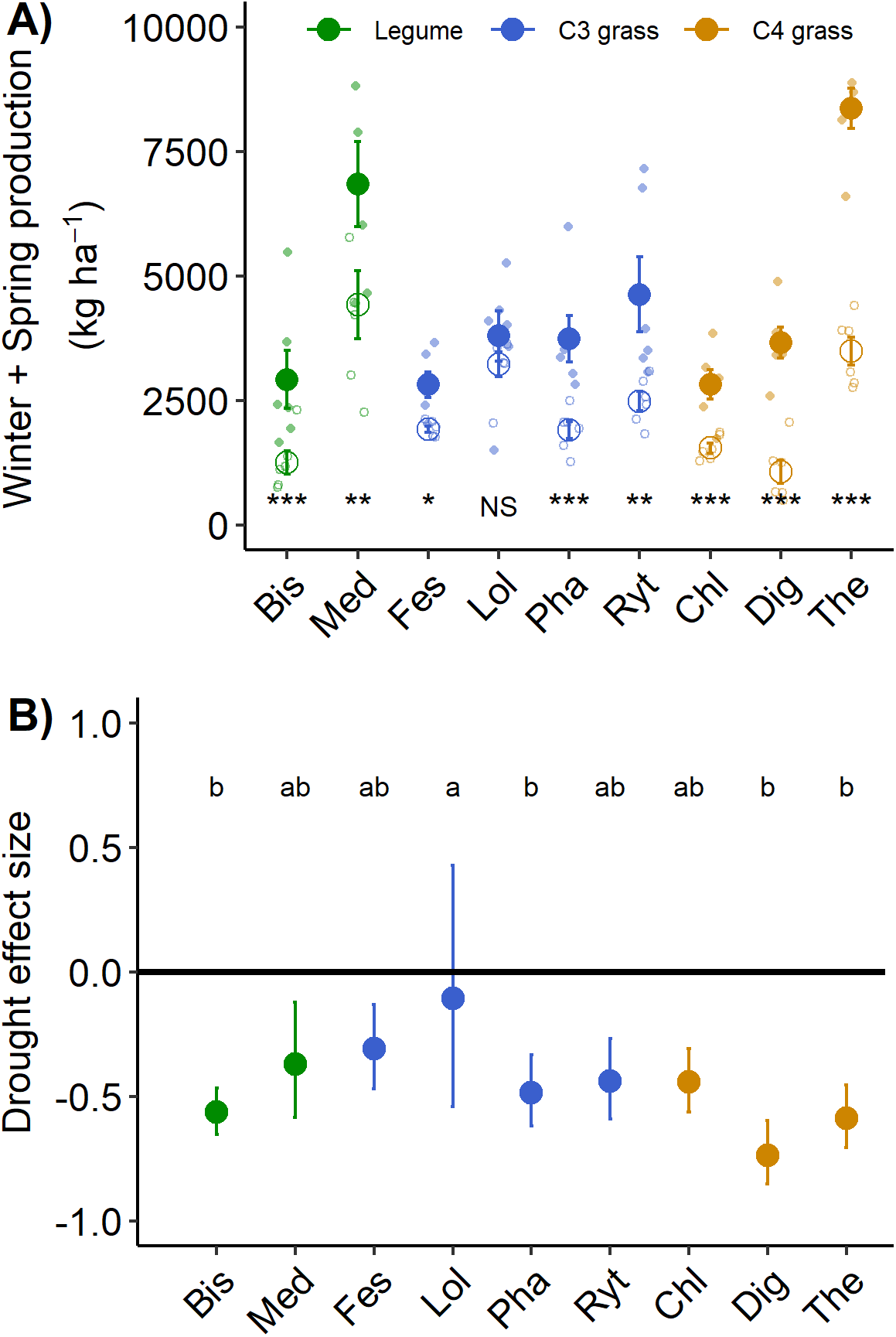
A) Aboveground production summed for all harvests during the 6-month winter/spring drought period. Large points shown are means ± 1 standard error (control = solid symbol, droughted = open symbol) and opaque points show species level variability in biomass. Significant pairwise comparisons for the effect of drought treatment are indicated as follows: NS= not significant, * *P* < 0.05, ** *P* = <0.01, *** *P* = < 0.001. B) drought effect size (log response ratio of drought vs control production during the 6-month drought treatment period). Values for panel B are mean values with 95% confidence intervals and same letter designations indicate non-significant differences among plantings. Abbreviations are as follows: *Biserrula* (Bis), *Chloris* (Chl), *Digitaria* (Dig), *Festuca* (Fes), *Lolium* (Lol), *Medicago* (Med), *Phalaris* (Pha), *Rytidosperma* (Ryt), *Themeda* (The).

**Table 2.** Linear mixed effects model output for the effects of drought on aboveground productivity of nine pasture and rangeland species during the 6-month winter/spring period.

| Response | Fixed Effects | F value | P value | R<sup>2</sup>m<sup>#</sup> | R<sup>2</sup>c<sup>\$</sup> |
| --- | --- | --- | --- | --- | --- |
| Biomass | Drought | 131.8<sub>1,23</sub> | <0.01 | 0.75 | 0.78 |
|  | Species | 24.0<sub>8,69</sub> | < 0.01 |  |  |
|  | Drought x Species | 4.2<sub>8,69</sub> | < 0.01 |  |  |
| Biomass | Drought | 114.8<sub>1,23</sub> | <0.01 | 0.27 | 0.78 |
|  | FG<sup>*</sup> | 0.0<sub>2,6</sub> | 0.97 |  |  |
|  | Drought x FG | 6.4<sub>2,84</sub> | <0.01 |  |  |

*All biomass data were ln transformed to meet assumptions of constant variance. F value subscripts indicate degrees of freedom, \*FG refers to functional group, <sup>#</sup>R<sup>2</sup>m indicates marginal error associated with linear model fixed effects, <sup>\$</sup>R<sup>2</sup>c indicates conditional error or the total variation described by the full model including nested random effects (plots within a shelter).*

The drought effect size, quantified as the log response ratio between control and droughted subplots, varied among species (Figure 4B) with the greatest reductions being shown by two of the C_4_ grasses (*Digitaria:* 74% and *Themeda:* 59% yield reduction). The two legume species showed intermediate reductions (*Biserrula*: 56% and *Medicago*: 37%), while C_3_ grasses were the most variable group (*Lolium*: 11% to *Phalaris*: 48% reduction). All plant functional groups were negatively impacted by drought, with C_4_ grasses experiencing significantly greater loss of productivity than C_3_ grasses or legumes (Figure 5; Table 2).

**Figure 5.**
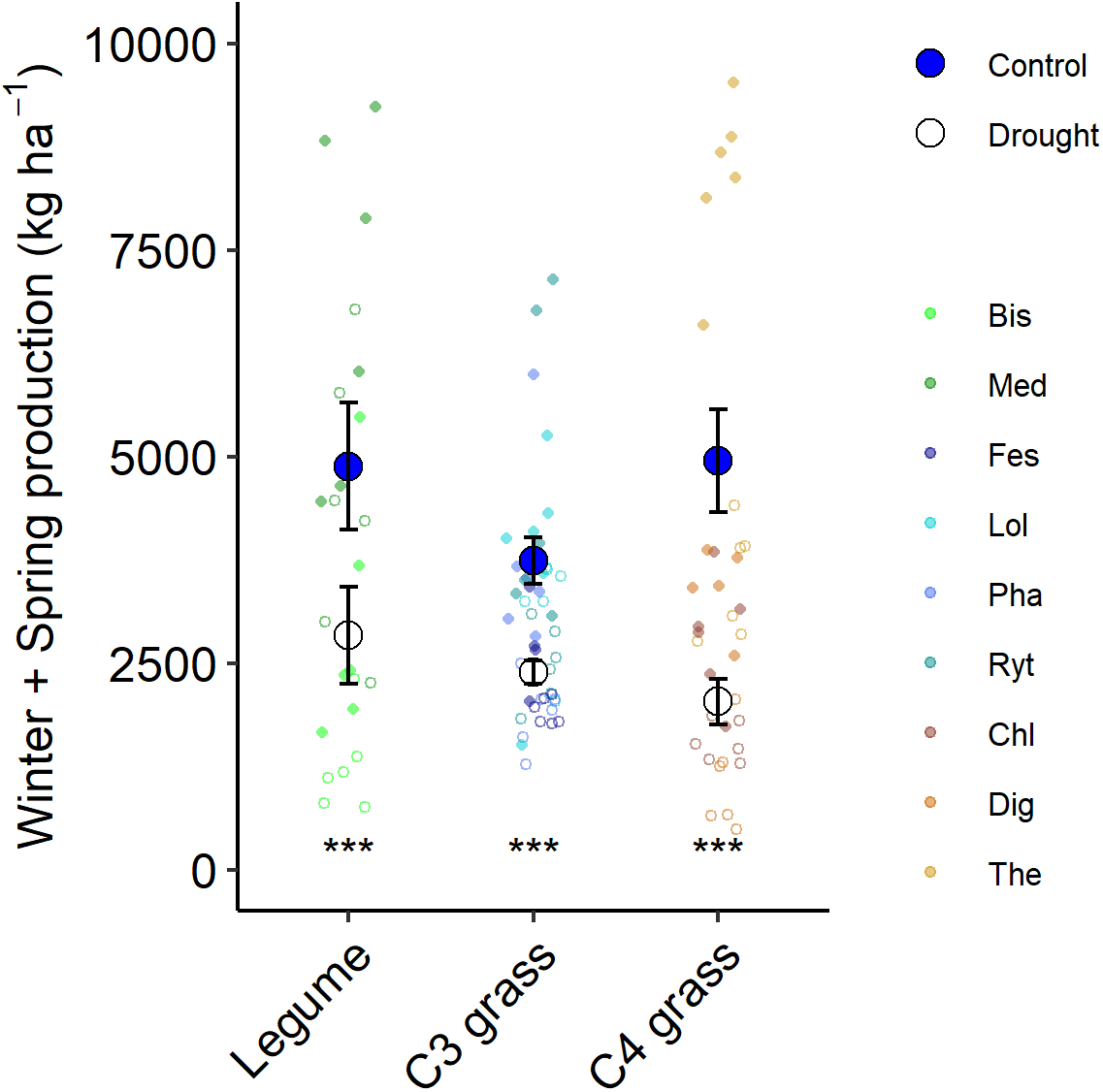
Aboveground production summed for all harvests during the 6-month winter/spring drought period by functional group. Large points shown are means ± 1 standard error (control = solid symbol, droughted = open symbol) and opaque points show subplot level variability in biomass for each species within the functional group. Significant pairwise comparisons for the effect of drought within a functional group are indicated matching Fig 4.

### 3.3 Effects of warming and drought-warming interactions on aboveground biomass production

Warming during the 6-month winter/spring period generally reduced productivity (Table 3). While there was a broadly consistent response to the combination of drought and warming (Table 3), the absolute magnitude of productivity decline differed between the two species (Figure 6A). Warming resulted in a significant decline in winter/spring productivity in *Festuca* that was greater under drought (control: 18 %, drought: 31% reduction; Figure 6B, C). *Medicago* swards were not significantly affected by warming under either precipitation treatment (Figure 6C). Additionally, while *Medicago* swards that experienced ambient temperatures were negatively impacted by drought, productivity of swards that were exposed to both warming and drought did not differ significantly from those receiving the control precipitation regime (Figure 6B). This indicates that warming slightly reduces the negative effects of drought for *Medicago* (Figure 6A, C), although interactions between warming and drought were not significant for either species (Figure 6D).

**Figure 6.**
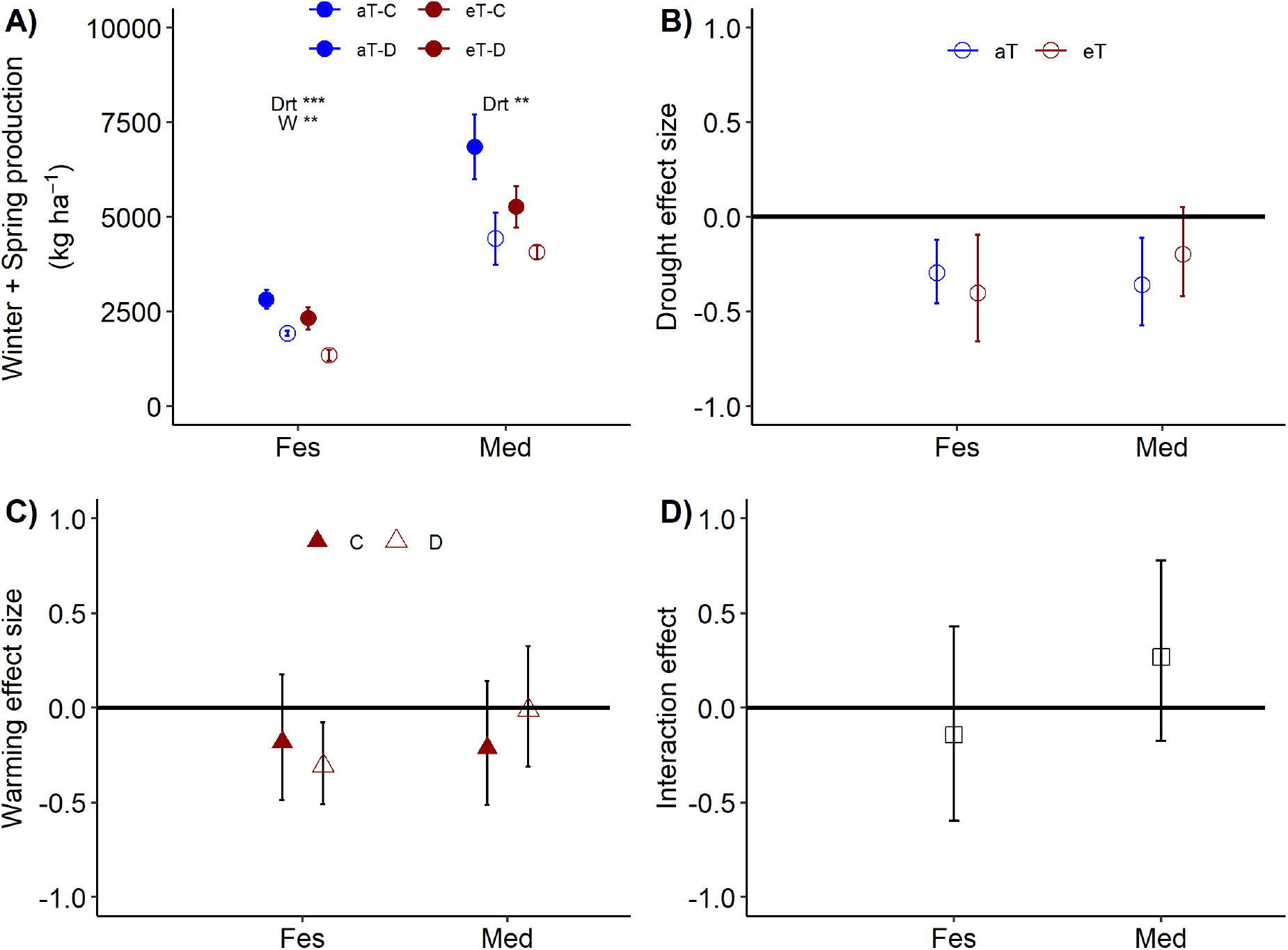
Aboveground production for *Festuca* and *Medicago* exposed to the combined effects of drought (control: C and droughted: D) and warming (ambient: aT and elevated: eT) treatments during A) the 6-month winter/spring drought period. Treatment effect sizes during the winter/spring drought period for B) drought under ambient (also shown in Fig 4) and elevated temperatures, C) warming, under control and droughted conditions, and D) the effect of warming on biomass responses to drought (for panel D, positive values indicate a reduction in drought impact under warming). Values in panel A are means ± 1 SE; while panels B-D are means and 95% confidence intervals. Spp. abbreviations and significance levels follow Fig. 4.

**Table 3.** Linear mixed effects model output for the combined effects of drought and warming on aboveground productivity of both *Medicago sativa* and *Festuca arundinacea* during the 6-month winter/spring period.

| Response | Fixed Effects | F value | P value | R<sup>2</sup>m<sup>#</sup> | R<sup>2</sup>c<sup>§</sup> |
| --- | --- | --- | --- | --- | --- |
| Biomass | Drought | 23.2<sub>1,15</sub> | <0.01 | 0.79 | 0.86 |
|  | Warming | 6.9<sub>1,15</sub> | 0.02 |  |  |
|  | Species | 208.3<sub>1,20</sub> | <0.01 |  |  |
|  | D x W | 0.0<sub>1,15</sub> | 0.86 |  |  |
|  | D x S | 0.7<sub>1,20</sub> | 0.42 |  |  |
|  | W x S | 1.6<sub>1,20</sub> | 0.22 |  |  |
|  | D x W x S | 2.5<sub>1,20</sub> | 0.13 |  |  |

*Biomass data were ln transformed to meet assumptions of constant variance, F value subscripts indicate degrees of freedom, <sup>#</sup>R<sup>2</sup>m indicates marginal error associated with linear model fixed effects, <sup>§</sup>R<sup>2</sup>c indicates conditional error or the total variation described by the full model including nested random effects (plots within a shelter).*

## 4 Discussion

Projections of future climate change, including increases in the frequency and magnitude of extreme climate events, are likely to disrupt ecosystem functioning (Jentsch and Beierkuhnlein, 2008; Knapp et al., 2008; Nijp et al., 2015), with important consequences for the productivity of pastures and rangelands (Jiménez et al., 2011; Godde et al., 2020). We found that eight of the nine C_3_ and C_4_ species exposed to extreme winter/spring drought experienced significant reductions in cool season productivity, with 45% declines on average among species and losses of up to 74%, relative to controls. Despite large species differences in drought sensitivity, we did find evidence for functional group-specific responses between C_4_ grasses, C_3_ grasses, and legumes, such that C_4_ grasses had the greatest response to drought. Furthermore, we found no evidence of a positive effect of warming on the productivity of two temperate species’ (*Festuca* and *Medicago*) and the combination of warming and drought resulted in the greatest biomass declines, which were either additive (*Festuca*) or less-than-additive (*Medicago*). Taken together, these results demonstrate the utility of evaluating responses of a range of species to single and compound climate extremes for improved forecasts of future grassland vulnerability to climate change.

### 4.1 Productivity responses to winter/spring drought

Decades of research focused on examining grassland responses to drought have emphasized the importance of regional climate context (Heisler-White et al., 2009; Knapp et al., 2017; Slette et al., 2019) including rainfall seasonality (Padrón et al., 2020). The timing of drought in relation to plant growth is a key factor influencing species’ responses to changing rainfall regimes, with evidence that shifts in rainfall seasonality can be more important than changes in the amount of rainfall (Belovsky and Slade, 2020). Cool season rainfall has been found to drive soil water storage and long-term patterns of productivity in many grasslands and regions with year-round patterns of plant growth (Derner et al., 2020). Furthermore, there is evidence that winter precipitation can offset the effects of spring/summer drought (Fry et al., 2013). Locally in South-eastern Australia, projections for increased severity of winter/spring drought (CSIRO, 2020) are likely to have large impacts on the seasonality of pasture and rangeland productivity. Similar shifts in precipitation may have far reaching implications for decisions on stocking densities and associated livestock production around the world (Derner et al., 2020).

Changes in the seasonality of precipitation and the timing of when drought occurs with a growing season, is particularly important in ecosystems that comprise a mixture of C_3_ and C_4_ plant species with distinct growth phenologies. In our warm temperate/subtropical region, we predicted that cool-season drought would more negatively impact C_3_ species due, in part, to the timing of high soil water deficits during their period of active growth as well as differences in water use efficiency. We found that while some individual C_3_ species (e.g. *Festuca*) experienced large growth reductions in response to cool-season drought, as a group, C_4_ species were more negatively affected than their temperate C_3_ counterparts. This finding has numerous implications for the sustainability of forage production, particularly in systems where growth occurs year-round due to a lack of cold temperature constraints. Firstly, sustained rainfall and soil water deficits during this period will have direct impacts on productivity of temperate species that are active in winter and early spring, contributing to winter feed-gaps. Secondly, the accumulated soil water deficit during periods of winter and early spring drought can have negative impacts on the spring growth of warm-season (C_4_) grasses, as seen in our study and those of others in the USA (Prevéy and Seastedt, 2014; Arredondo et al., 2016). If this response is consistent across C_4_ grass species, management recommendations that promote switching to more heat-tolerant C_4_ species to accommodate increased temperatures in traditionally temperate/sub-tropical cropping regions (Johnston, 1996) may not result in yield gains during the spring and early summer.

Legumes had the most consistent drought response, although broad generalizations are limited with results from only two species. Compared with other species in this study, *Medicago* experienced only modest impacts of drought on aboveground production. This may have been a consequence of deep tap roots (Li et al., 2012; Nie et al., 2015), allowing access to deep soil water to sustain growth during dry periods. Deep rooting strategies can also facilitate shifts in moisture depth profiles due to hydraulic lift (Raza et al., 2013), thereby reducing drought effects at the sward level (Liste and White, 2008; Pang et al., 2013). *Biserrula*, also a deep-rooted legume (Loi et al., 2005; Haling et al., 2016), experienced greater productivity losses in response to drought, compared to Medicago. However, unlike *Medicago*, a perennial whose swards develop over multiple years (Li et al., 2012), *Biserrula* is an annual species, regenerating from seed early in spring each year. These life history differences that influence growth seasonality along with the significant effect of winter/spring drought on productivity of the tropical (C_4_) grasses, highlight the importance of drought timing in relation to species’ growth phenology (Wilcox et al., 2017; Yao et al., 2019), which is at least partially reflected in their functional group classifications.

### 4.2 Plant responses to warming and drought × warming interactions

Gradual warming associated with climate change is predicted to affect ecosystems through a variety of mechanisms, including direct impacts of increased air and soil temperature on species’ physiology and indirect impacts on soil water content via increases in evaporation or transpiration (Deutsch et al., 2011). Many temperate and cold-climate systems report gains in productivity associated with warming (Bloor et al., 2010; Wang et al., 2012), due to reduced exposure to growth-limiting cold temperatures (Mori et al., 2014; Naudts et al., 2011; Reyer et al., 2013) or greater nutrient availability resulting from increased microbial activity (Bloor et al., 2010; Dellar et al., 2018). The beneficial impacts of warming on grassland productivity have been reported both globally (Gao et al., 2016) and locally (Cullen et al., 2009). In other systems, however, warming can shift temperatures beyond critical physiological thresholds, resulting in reduced growth or even tissue die-back (Bastos et al., 2014; Cremonese et al., 2017). In our sub-tropical system, we found no increase in productivity associated with cool-season warming for either pasture species. Rather, we found a significant overall decline in productivity in response to elevated temperature. This is despite an increase in winter growing degree days and a reduction in frost exposure, changes that are expected to increase winter and early spring growth (Chang et al., 2017; Piao et al., 2019). It is likely, therefore, that lower productivity was associated with supra-optimal temperatures for these species during spring and/or warming-associated reductions in water availability. In addition to these productivity changes, changes to the nutritional quality of pastures can be anticipated and will impact livestock production under future, more extreme climate conditions (Catunda et al., 2021).

*Festuca* and *Medicago* are widely planted across Europe, North and South America, Australasia and Africa (Gibson and Newman, 2001; Ghaleb et al., 2021), contributing to the pasture feed-base that underpins global livestock production. These temperate species have optimum temperatures for photosynthesis in the region of 20-29°C (*Festuca;* (Sasaki et al., 2002; Sinclair et al., 2007; Jacob et al., 2020) and 15-30°C (*Medicago;* Al-hamdani and Todd, 1990; Jacob et al., 2020). Whilst winter warming is, therefore, likely to stimulate gross photosynthetic rates, spring temperatures were regularly above these thresholds, especially in the later part of the season where daily maxima of over 45°C were recorded in warmed plots.

The exceedance of thermal optima, along with increased respiratory carbon losses at warmer temperatures (Heskel et al., 2016; Chandregowda, 2021) may explain observed productivity declines in response to warming. There was, however, also evidence of small reductions in soil water content associated with the warming treatment, particularly for *Festuca.* Given the generally low levels of soil water availability in these well-drained sandy soils, this increased soil moisture stress may have contributed to the large decline in aboveground productivity observed in warmed plots for this species. Although higher temperatures are generally associated with increased productivity in northern hemisphere grasslands (Craine et al., 2012), warming can have both positive (winter) and negative (summer) impacts, depending on ambient temperature (Kreyling et al., 2019). Contrary to expectations, our findings indicate that perceived benefits of winter and spring warming may not be realized under field conditions, especially where ambient levels of soil water availability are sub-optimal for growth. Similar negative relationships between grassland productivity and cool season temperatures have also been reported elsewhere from long term survey data (Wu et al., 2021).

High temperatures and drought are strongly coupled (Seneviratne et al., 2010) and their co-occurrence can exacerbate soil water deficits as a result of evaporation from surface soils and increased requirements for transpirational cooling (Ciais et al., 2005; Kirschbaum and McMillan, 2018). However, the ecological impacts of these co-occurring stressors depend on the physiological thermal optima and drought adaptation strategies of individual species, with additive, greater-than-additive or less-than-additive responses all reported (Zavalloni et al., 2008; Wu et al., 2011; Dreesen et al., 2012; Yu et al., 2012; De Boeck et al., 2016). In our study both species exposed to warming alongside drought experienced the greatest productivity declines in this combined treatment, with effects being either additive (*Festuca*) or less-than-additive (*Medicago*) such that the effects of winter-spring drought were lower reduced under continuous warming. An exacerbation of drought effects at higher temperatures has been reported across biomes and plant functional groups, often associated with reductions in soil moisture (Adams et al., 2009; Orsenigo et al., 2014; De Boeck et al., 2016), such as we found for *Festuca*. This connection between temperature and water availability is likely to amplify the intensity of ecological drought under future climates (Dai, 2011; IPCC, 2021).

Alternatively, temperature-driven reduction in available water has the potential to provide drought-priming effects, through ‘stress-memory’, that reduce impacts of subsequent water stress (Zavalloni et al., 2008). The less-than-additive productivity response in *Medicago* exposed to the combined warming and drought treatment suggests the possibility of plant acclimation to water stress, following prolonged exposure to low-severity droughts associated with the warming treatment (Schwinning et al., 2004; Walter et al., 2013; Backhaus et al., 2014). This acclimation may be important for the persistence and profitability of this pasture species in the future. Similar less-than-additive temperature and drought interactions have been observed in temperate grasslands and more generally across major biomes in meta-analyses (De Boeck et al., 2011; Wu et al., 2011; Song et al., 2019). Importantly, our results align with model predictions of productivity declines in Australian rangelands in response to moderate (+3 °C) warming combined with reduced rainfall (McKeon et al., 2009).

Many studies have highlighted the importance of a species’ persistence in drought and high temperatures (Culvenor et al., 2016), and this is especially true for perennial grasses and legumes in managed grasslands (Norton et al., 2009). Further work investigating the mechanisms underpinning species’ responses will help refine predictions about the impacts of multiple, co-occurring changes under future climates. This study provides important, new experimental, field-based data on the effects of extreme drought on a wide variety of pasture species and two native grasses, and, for a sub-set of species, in combination with continuous warming. These data complement information from modelling studies (Kaine and Tozer, 2005; Cullen et al., 2009) and comparisons across precipitation gradients (Clark et al., 2016) to quantify impacts of future, more extreme rainfall regimes on the productivity of the pasture and rangeland feed base that underpins livestock grazing in many parts of the world (Godde et al., 2020). Work is underway to characterise relationships between productivity losses and plant traits and strategies relating to the acquisition and use of water (root traits, plant-microbial interactions, plant hydraulics) and the allocation of carbon (above-versus belowground, root and crown carbohydrate storage), to determine the mechanisms responsible for the observed species’ differences to drought and warming. This information will be key to extrapolating findings from this study to a wider range of locations and grassland species, including many of international relevance in production systems across the globe.

## 5 Conclusions

This paper introduces a new experimental facility used to simulate future, more extreme climates, under field conditions. We found consistent negative effects of severe winter/spring drought on seven pasture species and two native Australian rangeland grasses, highlighting the challenges associated with future climate risk management for livestock production systems. Strong reductions in cool season productivity for all functional groups highlight potential climate constraints on winter forage availability, but also species’ persistence throughout the warmer summer months. Furthermore, the expanded use of tropical C_4_ grasses to mitigate declines in temperate C_3_ species with rising global temperatures may lead to lower cool-season productivity due to the relatively high seasonal drought sensitivity of the C_4_ grasses examined in this study. Trade-offs are therefore implicit in planting decisions aimed at enhancing pasture drought tolerance, given the increases in mean and maximum temperatures that are already being observed globally. Additionally, substantial productivity declines associated with warming, even in the cooler months, highlight the important role of temperature in altering ecosystem water balance and, potentially, carbon dynamics, suggesting limited benefits from future warming in warm temperate and sub-tropical systems. Selection of species and cultivars with the physiological and/or phenological traits that support sustained productivity under more extreme climate conditions will become increasingly important as climate change undermines the performance of traditional pasture and rangeland species.

## 6 Conflict of Interest

The authors declare that the research was conducted in the absence of any commercial or financial relationships that could be construed as a potential conflict of interest.

## 7 Author Contributions/Funding/Acknowledgements

The authors would like to acknowledge individuals whose efforts supported the data collection and site management for this work, including Angelica Vårhammar, Jackson Bridges-Parlett, Alexandra Boyd, Shania Therese Didier Serre, Samantha Weller, Jinyan Yang and Arjunan Krishnananthaselvan. We also gratefully acknowledge expert input to early project discussions from Cath Lescun, Tom Davison, Tom Dickson, Allen Newman, Warwick Badgery, Suzanne Boschma, Richard Eckard, Brendan Cullen and Alan Humphries, and to Heritage Seeds for providing seeds. This project was supported by funding from Meat and Livestock Australia’s Donor Company (P.PSH. 0793), Dairy Australia (C100002357) and Western Sydney University. Authors ACC and SAP wrote the paper with input from all co-authors on draft iterations; ACC conducted statistical analyses with input from BEM, HZ, JRP and KJF; SAP, EP, MGT, DTT, CVMB, BA, JRP, JMP, CAM, BDM, YC, ICA designed the field experiment with input from ACC, HZ and KJF; ACC, HZ, KJF, KLMC, MC, CI, VJ, GWK and AKP collected field samples.

## 8 Data Availability Statement

The datasets analysed for this study can be found in the Dryad Repository upon publication of this manuscript.

## Notes

### Competing Interest Statement

The authors have declared no competing interest.

### Summary of Updates

This version was created in response to reviewer comments from the originally posted document, and reflects a new interval of sampling period and emphasis on a subset of pastures from the initial manuscript. All future version and subsequent publication will use this modified dataset.

