## Supplemental Materials for "Pastures and Climate Extremes: Impacts of cool season warming and drought on the productivity of key pasture species in a field experiment"

**Churchill et al.**

**Supplemental Information 1**

**Table S1)** Soil properties (0 – 10 cm depth) at PACE field site (September 2017)

| **Soil property** | **Measurement (units)** |
|---|---|
| Texture | 81.3% sand |
| | 7.5% silt |
| | 11.2% clay |
| Soil type | Loamy sand |
| Bulk density | 1.2g $cm^{-3}$ |
| Organic carbon (OC) | 1.0 ± 0.1 % |
| Organic matter OM | 1.8 ± 0.1% |
| pH ($CaCl_2$) | 4.9 ± 0.2 |
| pH (water) | 5.7 ± 0.2 |
| Nitrate -N | 42.4 ± 8.8 mg $kg^{-1}$ soil |
| Ammonium- N | 3.8 ± 1.1 mg $kg^{-1}$ soil |
| $PO_4$ (Bray P) | 25.7 ± 14.0 mg $kg^{-1}$ soil |
| $PO_4$ (Colwell P) | 39.0 ± 18. 4 mg $kg^{-1}$ soil |
| Phosphorus buffering index (PBI) | 30.9 ± 4.6 |
| Sulphur | 5.9 ± 0.6 mg $kg^{-1}$ |
| Exchangeable Aluminium | 2.0 ± 1.0 mmol $kg^{-1}$ |
| Exchangeable Calcium | 23.0 ± 4.0 mmol $kg^{-1}$ |
| Exchangeable Magnesium | 6.0 ± 1.0 mmol $kg^{-1}$ |
| Exchangeable Potassium | 6.0 ± 1.0 mmol $kg^{-1}$ |
| Exchangeable Sodium | 0.7 ± 0.1 mmol $kg^{-1}$ |
| Electrical conductivity (Salinity) | 0.10 ± 0.01 dS $m^{-1}$ |
| Boron | 0.36 ± 0.04 mg $kg^{-1}$ |
| Copper | 12.3 ± 1.4 mg $kg^{-1}$ |
| Iron | 41.6 ± 6.6 mg $kg^{-1}$ |
| Manganese | 71.2 ± 5.4 mg $kg^{-1}$ |
| Zinc | 2.2 ± 0.5 mg $kg^{-1}$ |

**Table S2)** Planting establishment timeline and cultivar information

| Species Name* | Cultivar | Seeding initiated | Tube stock planted | Planting established Re-seeding dates | Pilot year sward yield† 06/2018-05/2019 (kg ha$^{-1}$ yr$^{-1}$) |
|---|---|---|---|---|---|
| *Biserrula pelecinus* | Casbah | 04/2018 | NA | 08/2018 | 1776 ± 367 |
| *Medicago sativa* | SARDI 7 series2 | 08/2017 | NA | 08/2018 | 11201 ± 1225 |
| *Festuca arundinacea* | Quantum II MaxP | 08/2017 | NA | <05/2018 | 5074 ± 508 |
| *Lolium perenne* | Kidman | 04/2018 | NA | 08/2018 | 2348 ± 327 |
| *Phalaris aquatica* | Holdfast GT | 08/2017 | NA | 10/2018 | 5801 ± 680 |
| *Rytidosperma caespitosum* | Evans | NA | 03/2018 | 10/2018 | 7790 ± 912 |
| *Chloris gayana* | Katambora | 10/2017 | NA | <05/2018 | 9664 ± 980 |
| *Digitaria eriantha* | Premier | 10/2017 | 04/2018 | <05/2018 | 10507 ± 1638 |
| *Themeda triandra* | Badgerys Creek, NSW | NA | 03/2018 | <05/2018 | 17384 ± 485 |

**All species are referenced by genera in text, †Sward yield reported for monoculture ambient temperature, control precipitation subplots*

**Table S3.** Locations of permanently installed soil moisture (TDR) and temperature probes in subplots at the PACE experimental facility

| Species | Treatment | Sensor | Depth |
|---|---|---|---|
| *Biserrula pelecinus* | Control, Drought | TDR | 0-15 cm |
| *Festuca arundinacea* | Drought *Warming combinations | TDR<br>Temperature | 0-15 cm<br>6-12 cm |
| *Lolium perenne* | Control, Drought | TDR | 0-15 cm |
| *Medicago sativa* | Drought * Warming combinations | TDR<br>Temperature | 0-15, 15-30 cm<br>6-12 cm |

**Table S4**. Mean and ± standard errors for surface temperature measurements (° C) determined by IR sensor thermography for each treatment plot type between June 1st and November 30th 2019

| Treatment | Surface Temperature |
|---|---|
| aT-C | 16.6 ± 0.1 *d* |
| aT-D | 17.6 ± 0.1 *c* |
| eT- C | 19.5 ± 0.02 *b* |
| eT-D | 20.5 ± 0.02 *a* |

*Same letters indicate non-significant differences among pairwise drought (control: C, drought: D) and warming (ambient: aT, elevated: eT) treatments*

**Table S5.** Total number of days between June 1st and November 30th, 2019, where treatment plots exceeded specified surface temperature thresholds, monitored at the 4 x 4 m plot level using IR sensors.

| | **aT-C** | **aT-D** | **eT-C** | **eT-D** |
|---|---|---|---|---|
| Days < 0 °C | 1 | 0 | 0 | 0 |
| Days < 5 °C | 45 | 42 | 15 | 9 |
| Days > 35 °C | 47 | 66 | 57 | 79 |
| Days > 40 °C | 21 | 45 | 37 | 56 |
| Days > 45 °C | 6 | 23 | 14 | 37 |

**Commented [AC1]:** @Kate Fuller- Can you check my calculations here compared with your code?

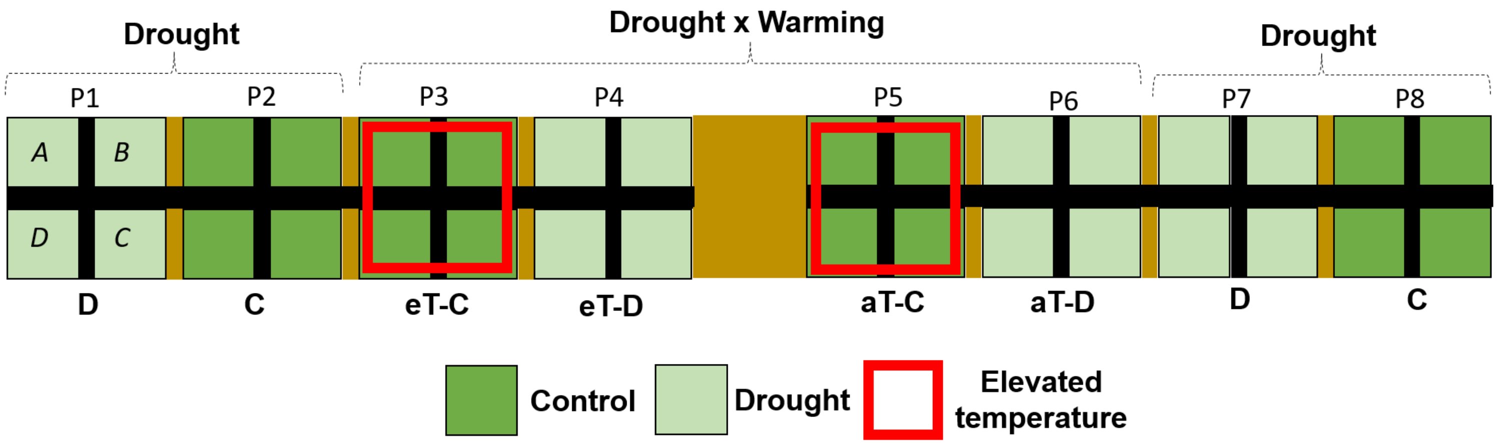


**Figure S1**. Schematic showing example shelter configuration of treatment plots (P1-8) and nested planting specific subplots (*A-D*) at the PAstures and Climate Extremes field facility. Experimental effects of warming (ambient: aT, elevated: eT) and drought (control: C, drought: D) treatments.

The PAstures and Climate Extremes (PACE) facility comprised six rainout shelters (S1-6), laid out as in **Figure S1**. Drought and warming treatments were applied at the plot level (P1-8, 4 x 4 m), each with four 2 x 2 m subplots nested within (*A-D*). Plantings of one of twelve different pasture species or two species mixtures were assigned at the subplot level. All plantings exposed to both drought and warming treatments (*Festuca*, *Medicago*, *Phalaris*/*Trifolium* and *Themeda*/*Rytidosperna*) were grouped together and randomly assigned subplot locations within plots 3-6, while plantings exposed to only drought treatments were grouped into two pairs of plots – P1+P2 and P7+P8, (with one droughted and one control plot in each pair), and species assigned randomly to subplots within each pair.

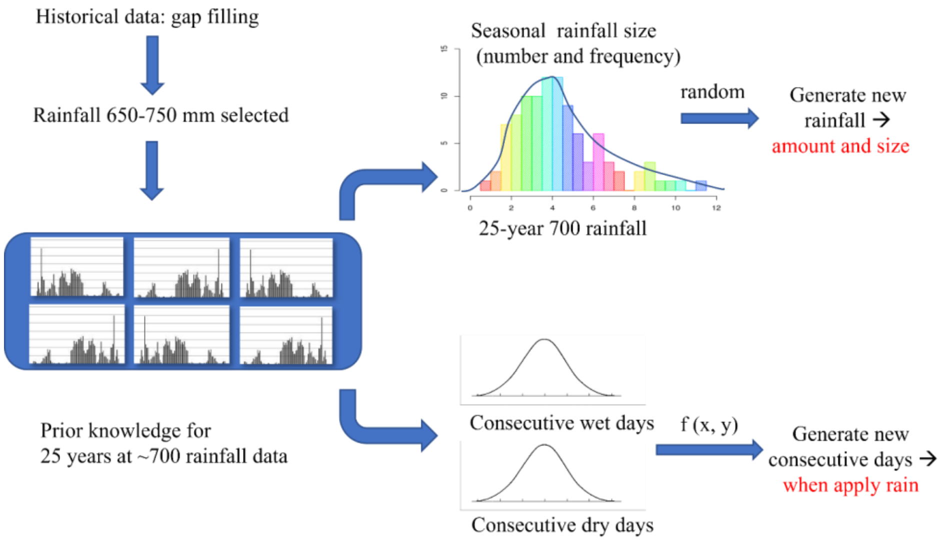


**Figure S2**. Detailed steps for generating the ambient control rainfall regime based on long-term historical rainfall data

We generated the control rainfall regime based on historical, local site data for Richmond, NSW. This comprised 128 years of daily rainfall records supplied by the Australian Bureau of Meteorology for the period 1885-2013. We used a gap-filling method to fill in missing data and selected years in which annual rainfall ranged from 650 to 750 mm, resulting in a database containing daily rainfall data for 25 years. This was used to summarise the statistical characteristics of rainfall patterns, by season, in terms of (1) mean total rainfall amount, event size distribution (small, <5 mm; medium, 5-30 mm; big, > 30 mm) and specific number of events in each size category; (2) the consecutive number of wet and dry days (continuous rain or dry days). This characterisation was used to randomly generate a control rainfall pattern for each season. Detailed steps are outlined in **Figure S2**. Rainfall was then applied as scheduled irrigation events with the exact volume controlled using a water meter (T-Probe, Elster).

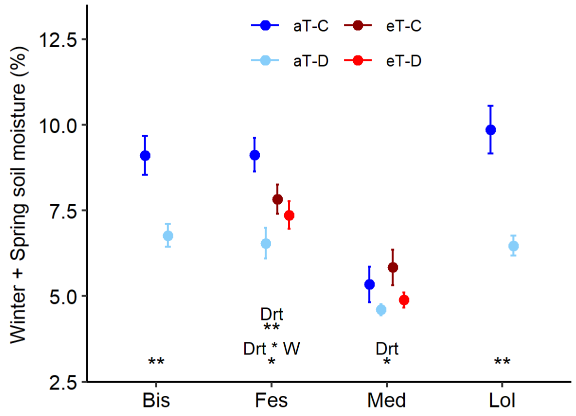


**Figure S3**. Mean soil volumetric water contents (%) among the four continuously-monitored species during the 6-month winter/spring drought period (June 1st-Nov 30th 2019). Points shown are means ± 1 SE, and all species abbreviations match Fig 3. Treatments are drought (Drt; control (C) and droughted (D) plots) and warming (W; ambient: aT, elevated: eT). Symbols (NS- not significant, *: $P \leq 0.05$. ** $P$: $\leq 0.01$, ***: $P \leq 0.001$) indicate statistical differences in mean soil volumetric water content among treatments for a species.

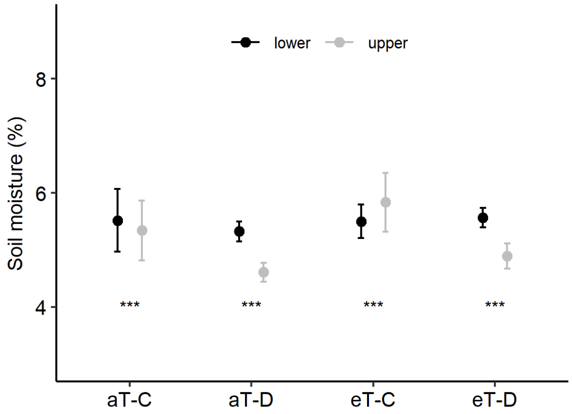


**Figure S4**. Soil volumetric water content monitored at two depths (upper: 0-15 cm and lower:15-30 cm) in *Medicago* subplots during the 6-month winter/spring drought period. Points shown are means ± 1 SE. Symbols (NS- not significant, *: $P \leq 0.05$. ** $P$: ≤ 0.01, ***: $P \leq 0.001$) indicate statistical differences in mean soil volumetric water content between lower and upper soil depths within each drought (Control: C and Droughted: D) and warming (ambient air temperature: aT and elevated air temperature: eT) treatment combination.

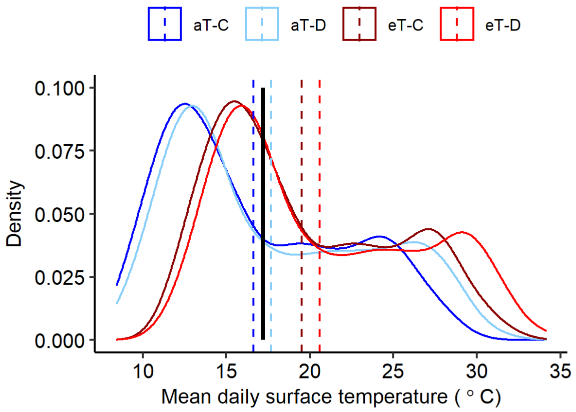


**Figure S5**. Density plot of daily mean surface temperatures for plants and soils in plots associated with IR lamp warming treatments among ambient (aT) and elevated (eT) treatments for both control (C) and droughted (D) plots throughout the year (June 1st 2019- November 30th, 2019). Dashed lines indicate treatment mean annual surface temperature, and the solid black line indicates long-term (1980-2010) mean annual air temperature in Richmond, NSW Australia.

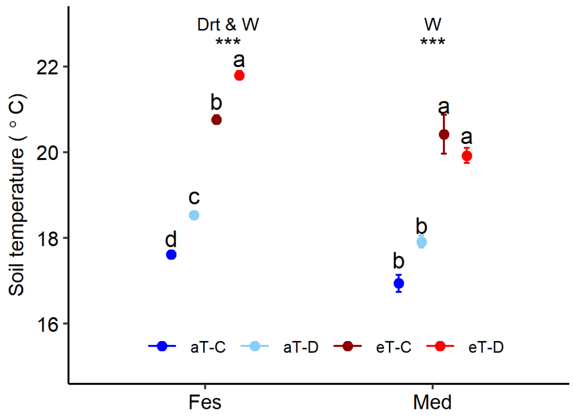


**Figure S6**. Average soil temperature measured among drought and warming treatments at 6-12 cm depth in *Festuca* (Fes) and *Medicago* (Med) plantings during the 6-month winter/spring drought period. Points shown are means ± 1 SE, symbols indicate significant effects of treatments within a species, while same letters indicate non-significant differences among treatment combinations within a species. Treatments are drought (Drt; control (C) and droughted (D) plots) and warming (W; ambient: aT, elevated: eT).

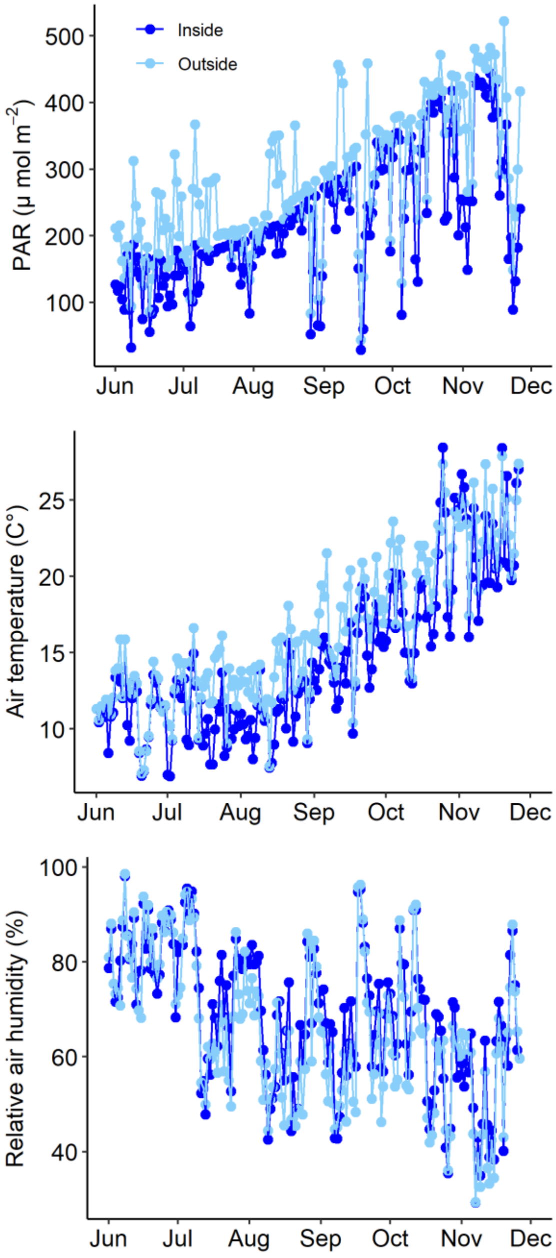


**Figure S7.** Average daily measurements for A) PAR, B) air temperature, and C) relative air humidity between sensors inside shelters and outside shelters to examine the impact of the PACE shelter infrastructure on environmental conditions.
